# A survey of transcriptomic datasets identifies ABA-responsive factors as regulators of photomorphogenesis in *Arabidopsis*

**DOI:** 10.1101/2022.07.24.501316

**Authors:** Cássia Fernanda Stafen, Iara Souza, Ben Hur de Oliveira, Luísa Abruzzi de Oliveira-Busatto, Rodrigo Juliani Siqueira Dalmolin, Oscar Lorenzo Sánchez, Felipe dos Santos Maraschin

## Abstract

Following germination, seedlings grown in light show a photomorphogenic development with open and green cotyledons and a robust root system. The light perception by the photoreceptors activate autotrophic photosynthetic metabolism to sustain growth of the whole plant. Several studies have evaluated transcriptional responses to light signals. Nevertheless, evaluating a single source experiment might bias the identificationof general, reproducible light responses. In order to identify widespread light-dependent signaling events that control early seedling photomorphogenesis we performed a survey comparing commonly regulated genes in transcriptomic public datasets derived from etiolated seedlings exposed to short light treatments. By compiling commonly regulated genes from different datasets, we obtained broadly representative regulated processes concerning general light transcriptional response. Our analysis shows that light primarly affects shoot gene expression promoting the assembly of photosynthetic machinery, signaling and redox responses. We observed that Transcriptograms allowed a better comparison among different experiments than DEseq analysis. We also identified that, transcriptional regulation of early light response is centered in the transcription factor ABA-Insensitive5 (ABI5) along with other bZIP transcription factors suggesting a mechanism by which dark expressed transcription factors guide the activation of early photomorphogenic genes.

**Key Message:** Comparative analisys of seedling deetiolation transcriptomic datasets identified ABA-responsive bZIP transcription factors as central regulators of early photomorphogenesis

## Introduction

Light is one of the most important enviromental factors that controls plant growth and development. Plants have evolved multiple ways to perceive light-related changes, such as quantity, quality, period and direction, adapting to ajust their development accordingly (Mancini et al. 2016; Dong et al. 2017; Oh et al. 2019; Lacek et al. 2021). The signaling system consists of photoreceptors for blue/UV-A light (phototropins, cryptochromes, ‘Zeitlupes’), red/far-red (phytochromes), and UV-B (UVR8) (Yu, 2010; Arsovski et al. 2012;Tilbrook et al. 2013; Christie et al. 2015; Galvão and Fankhauser 2015; de Wit M, Keuskamp DH, Bongers FJ 2016; Yang et al. 2017; Pham et al. 2018). Photomorphogenic development is dependent on a large transcriptional reprogramming downstream of photoreceptor signaling. Photomophogenesis is actively repressed by many regulators such as the CONSTITUTIVE PHOTOMORPHOGENIC/DE-ETIOLATED/FUSCA (COP/DET/FUS) and PHYTOCRHOME INTERATING FACTOR (PIF) signaling modules (Chory 1993; Hellmann and Estelle 2002; Lau and Deng 2012; Xu et al. 2015). The main receptors that control photomorphogenesis are the Phytochromes (phy) and Cryptochromes (cry) that upon light activation promote degragation of PIFs and supression of COP1/SPA ubiquitin ligase complexes, allowing the accumulation of light response transcrition factors such as ELONGATED HYPOCOTYL 5 (HY5), ELONGATED HYPOCOTYL HOMOLOG (HYH), LONG AFTER FAR-RED LIGHT1 (LAF1), among others that trigger deetiolation (Lau and Deng 2012; Xu et al. 2015; Zhong et al. 2021).

*Arabidopsis* seedlings display contrasting developmental phenotypes when grown under light or in the darkness. In the light, the seedlings present a photomorphogenic development with green and open cotyledons, a short hypocotyl and a long primary root (Wei et al. 1994). In darkness, however, the skotomorphogenic development produces seedlings with closed cotyledons and an apical hook, a long hypocotyl and a short root (Pham et al. 2018). Under the ground, the seedling protects the apical meristem keeping closed cotyledons and stimulate hypocotyl elongation. When exposed to light, the cotyledons open, expand and start to perform photosynthesis allocating resources for root growth (Lau and Deng 2012). The shoot usually consists of the primary light responsive organ in the plant, but it also controls the developmental responses of the roots underground (Van Gelderen et al. 2018). However, the shoot-derived signals that promote root photomorphogenic growth are still under debate, such as sugars, phytohormones, the HY5 transcription factor among others (Marchant et al. 2002; Salisbury et al. 2007; Laxmi et al. 2008; Kircher and Schopfer 2012; Regnault et al. 2015; Chen et al. 2016; Yang et al. 2020; Bhagat et al. 2021). A large amount of transcriptomic data is available from Arabidopsis seedlings exposed to different light conditions. Although these datasets have been evaluated individually, a crossed comparison of these data may help to find common genes involved in light perception and signaling with higher confidence. In order to explore genes and pathways involved in early shoot light responses, we aimed for integrating expression profiling studies in *Arabidopsis* performing a survey in published public transcriptomic databases from light-treated young Arabidopsis etiolated seedlings. Our analysis allowed us to identify ABA-response factors as putative primary regulators of photomorphogenic gene expression linking dark-expressed transcription factors to guide the activation of early photomorphogenic genes.

### Materials and methods Plant materials

*Arabidopsis* Columbia (Col-0) was used as wild-type (WT), and the mutant *hy5* (SALK_056405C) is in Col-0 ecotype background and were obtained from the The European Arabidopsis Stock Centre (NASC, http://arabidopsis.info/), *abi5-7* (Oh et al. 2009a), *35ScMyc:ABI5, pABI5:ABI5-GUS* (Mallappa et al. 2008a) *abf3 (SALK_075836), and abf4 (SALK_069523)* (Fernando et al. 2018) were kindly donated by Dr. Dana F. Schroeder.

### Data Mining

For data collection, the NCBI GEO functional genomics repository was queried. Experiments under this platform were searched using keywords, “*Arabidopsis*”, “light” AND “shoot”. The microarrays selected were, GSE5617 (AtGenExpress database, http://bar.utoronto.ca/efp/cgi-bin/efpWeb.cgi) (Kilian et al. 2007; GSE29657 (Liu et al. 2012) as well as the RNA-seq GSE79576. These datasets were selected because of their similar growth conditions, similar age of seedlings and wild ecotype (Col-0). Supplementary Table 1 summarizes the experimental conditions of the data sets.

### Individual preprocessing and differential expression analysis of the datasets

For the microarray samples, the raw data was normalized with RMA method and expression values (Robust Multi-Array Average Expression Measure) were retrieved with affy R/Bioconductor package (Gautier et al. 2004). Differential expression analysis was performed using limma R/Bioconductor package. Genes were considered differentially expressed when p value _≤_ 0.001 with FDR adjustment (Ritchie et al. 2015). For the RNA-seq databases, the raw reads were aligned and annotated to *Arabidopsis thaliana* reference genome v10 using STAR aligner (Dobin et al. 2013). Counts from aligned reads were obtained with featureCounts (Liao et al. 2014) and normalized into log-2-counts-per-million. Differential expression analysis was performed using limma R/Bioconductor package protocol for RNA-seq experiments, considering only the differentially expressed genes with p value ≤ 0.001 with FDR adjustment (Ritchie et al. 2015).

### *Arabidopsis thaliana* transcriptogram

Protein-protein interaction data for *Arabidopsis thaliana* was obtained from the STRING v.10.5 database (Szklarczyk et al. 2017) using an associated cutoff score of 0.700. The Transcriptogramer v1.0 (Rybarczyk-Filho et al. 2011; da Silva et al. 2014) software was used for protein ordering. The Transcriptogramer package (Morais DA 2017) was applied to generate the transcriptograms of all datasets and their treatments, these were plotted as: for each position i of the indexes of ordering in the transcriptograms, the relative expression value assigned to such position will be given by

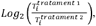

Where Ti is the average expression of the genes encoding for proteins belonging to the ith windowed range. The reading counts mapped to each individual gene determined the values of gene expression.The Monte Carlo sampling process allowed the localization of significant peaks and valleys in transcriptograms, in which random sets of permutations and sorting indexes were designed and used to randomly generate transcriptograms, from which zero distributions of peaks and lengths of valleys were inferred. These null distributions served as a basis for determining critical values for statistically significant peaks and valleys (p <0.001) and the number of permutations for each of the tests was established after convergence of the critical values. In order to perform this statistical analysis, internal scripts were used in the R programming environment. Functional enrichment was performed through the topGO package (Fisher test with Benjamini-Hochberg correction; Alexa and Rahnenfuhrer, 2016) and the GO annotations for the genes of the species studied were performed by the biomaRt package (Durinck et al. 2009) obtained from the ENSEMBL Plant database.

### GO terms funcional enrichment

The gene enrichment analysis on the significant peaks and/or valleys of the transcriptograms and DEseq was performed by aggregation ontology for the terms of biological processes (BP) using the website agriGO (v2.0) (Tian et al. 2017). The ReViGO (Supek et al. 2011) was used for remove redundant terms, calculate and summarize the list of GO terms according to the biological process.

### Transcription factor investigation

The list of DEGs and peaks/valleys at the intersection between the datasets GSE5617, GSE79576 and GSE29657 was evaluated for putative regulating transcription factors. Each of the four lists (DEseq up / down and Transcriptomic peaks/valleys) was analysed in TF2Network (http://bioinformatics.psb.ugent.be/webtools/TF2Network/) (Kulkarni et al. 2018) and the transcrption factors ranked with a Protein-DNA(PD) score ≥10% were selected for generating their interaction network with GeneMania plug-in (Mostafavi et al. 2008) in Cytoscape software (Shannon et al. 2003).

### Growth conditions for mutant phenotyping

Seeds were surface-sterilized and stratified at 4 °C for 2 days in complete darkness. Seedlings were grown at 21 °C ± 2 °C under completely darkness for 4 days and transferred to 1 and 4 hours of exposure white light illumination (94 _μ_mol m^−2^ s^−1^, white 6500k Led lamps). Plants were grown on half-strength sucrose-free Murashige and Skoog medium (Murashige and Skoog 1962) supplemented with 1.5% agar (w/v; KASVI) and 0.05% MES hydrate (w/v; Sigma-Aldrich, M8250), pH 5.7, on vertically oriented square plates. For dark grown plants, the plates were covered with aluminum foil. Similarly grown seedlings were used for RNA isolation. For root and hypocotyl length measurements, seedlings were growth for 14 days in LD condition (LD - Shoot light, Dark root, as Miotto et al. 2019), and the measurements were performed at 4, 7, 10 and 14 days. Organ length was measured with ImageJ (Fiji).

### Analysis by qRT□PCR

Total RNA was isolated from 4 days-old *Arabidopsis* seedlings grown as above using TRIzol™ Reagent (Thermo Fischer, #15596026). The cDNA was synthesized using M-MLV reverse transcriptase (Promega, # M5313) following the manufacturer’s instructions. qRT-PCR was performed in a StepOne(tm) Real-Time PCR System using Platinum Taq DNA Polymerase (Thermo Fischer, # 10966) according to the manufacturer’s protocol. Three independent biological samples were analyzed for each datapoint with three independent technical replicates each. The transcript levels were normalized against the reference gene AT3G18780, and relative expression was calculated by the ddCt method (Livak and Schmittgen, 2008) (Supplementary table 6).

### Chlorophyll quantification

Chlorophyll measurement was performed essentially as described (Holm et al. 2002). Briefly, 3-DAG seedlings harvested in darkness and after exposure to ligth for 1h, 4h, 6h, 8h and 24h were weighed, frozen in liquid nitrogen, and ground to a fine powder. Total chlorophyll was extracted with 80% acetone, and chlorophyll a and b contents were calculated using MacKinney’s specific absorption coefficients in which chlorophyll a = 12.7(A663) − 2.69(A645) and chlorophyll b = 22.9(A645) − 4.48(A663). The total specific chlorophyll content is expressed as micrograms of chlorophyll per gram of seedlings.

### Protochlorophyllide assay

The protochlorophyllide measurements were performed essentially as described (Job and Datta 2021). In total, 50 seedlings of were grown as above for 5-DAG in the dark (plates were covered in layers of aluminum foil). Seedlings were immediately frozen in liquid nitrogen and extracted with 90% acetone with 0.1 M NH4OH and incubated at 4°C for 1 h. Samples were centrifuged at 14000 g for 10 min and the supernatant was collected. Fluorescence spectra was measured with excitation at 440 nm and the emission spectra was recorded from 600 nm to 750 nm.

### GUS staining and microscopy analysis

Seedlings were fixed in 80% acetone for 20 minutes at -20ºC, washed 3 times in water and incubated overnight in GUS staining buffer [10 mM EDTA, 100 mM sodium phosphate (pH 7.0), 0.1% (v/v) Triton X-100, 1 mM K3Fe(CN)6, 1 mM K4Fe(CN)6, 1 mg/ml 5-bromo-4-chloro-3-indolyl-D-glucuronide] at 37 □. Subsequently, samples were washed in water once and cleared in 70% (v/v) ethanol at room temperature before imaging.

## Results

### Transcriptogram analysis of early and late light responses

We generated a transcriptogram considering a window of 17500 genes (Fig. 1a) and plotted the transcriptogram with all datasets (Fig. 1b), and further separated samples of early light responses (up to 1 hour after light exposure) (Fig. 1c) and late response (4 to 6 hours of light exposure) (Fig. 1d). Individual transcriptograms for each datasets are presented in Fig. S2. When overlaid, most transcriptograms showed a similar distribution profile of peaks and valleys (Fig. 1b), although with some clear amplitude variations. Most of peaks and valleys encompass GOs terms such as “cellular nitrogen compound metabolic process”, “response to light stimulus”, “photosynthesis”, “protein transport”, “protein import into nucleus”, “glutathione metabolic process ” (Fig. 1a, b). When we compared the light response profile over time, we could observe an increase in the amplitude of significant peaks and valleys in the late responses when compared to early responses, suggesting that the general expression profiles are maintained over time and are mostly quantitatively affected by prolonged light exposure.

**Figure 1.**
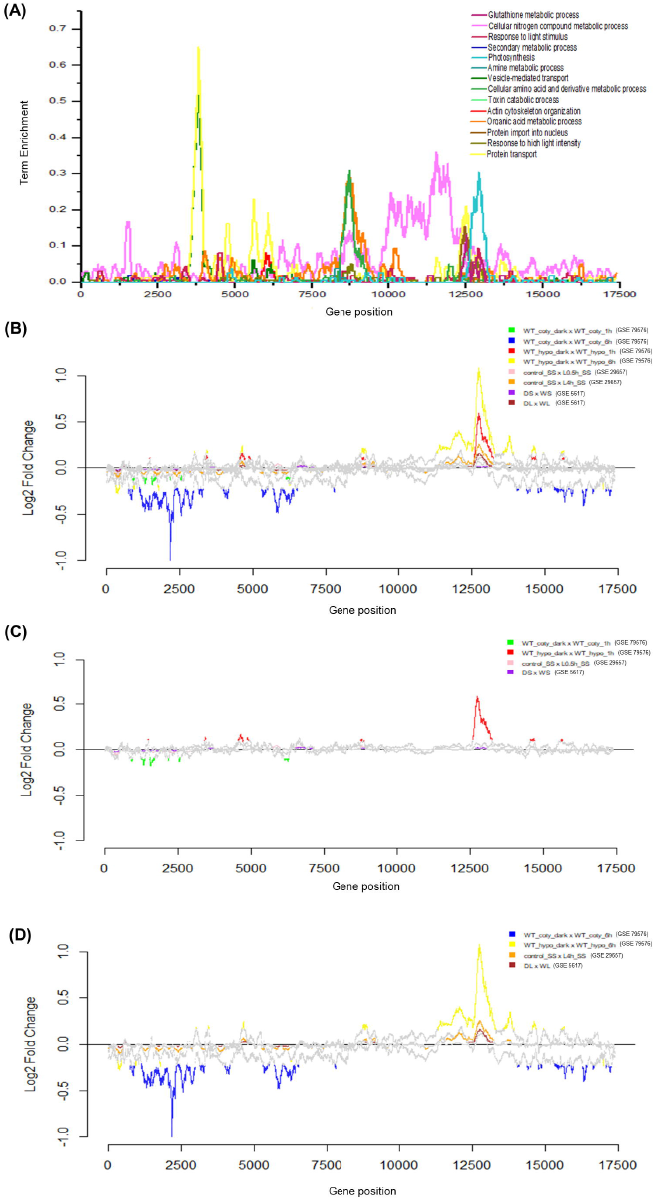
*Arabidopsis thaliana* transcriptogram comparison among three light response datasets. **a** *Arabidopsis thaliana* transcriptogram. The x-axis relative to gene position have been divided by the total number of proteins retrieved from STRING. Projection of Gene Ontology terms is color-coded. **b** Overlay of all transcriptograms for GSE5617, GSE29657 and GSE79576 datasets, **c** early responses GSE5617 (1 h WL), GSE29657 (0.5 h WL) and GSE79576 (1 h WL, coty - cotyledons and hypo - hypocotyls) and **d** late light responses GSE5617 (4 h WL), GSE29657 (4 h WL), and GSE79576 (6 h WL, coty - cotyledons and hypo - hypocotyls). Average transcriptograms signifcant peaks (up-regulation) or valleys (down-regulation) are represented by colors. Grey lines show non-signifcant regions

### Identification of common light regulated genes

In order to compare the similarity between the datasets, an intersection with the common modulated genes present in the significant peaks and valleys of all the transcriptograms as well as differentialy expressed according to DEseq was generated (Fig. 2). From the comparison of these datasets, 920 genes overlaped in significant peaks (Fig. 2a) whereas 619 genes overlapped in significant valleys (Fig. 2b). When we compared the same datasets by DESeq analysis we found an overlap of 214 up-regulated (Fig. 2c) an 456 down-regulated genes among the three datasets listed above (Fig. 2d). Based on these observations, we conclude that the transcriptogram is able to identify a larger set of regulated genes than DEseq allowing a broader analysis of physiological responses.

**Figure 2.**
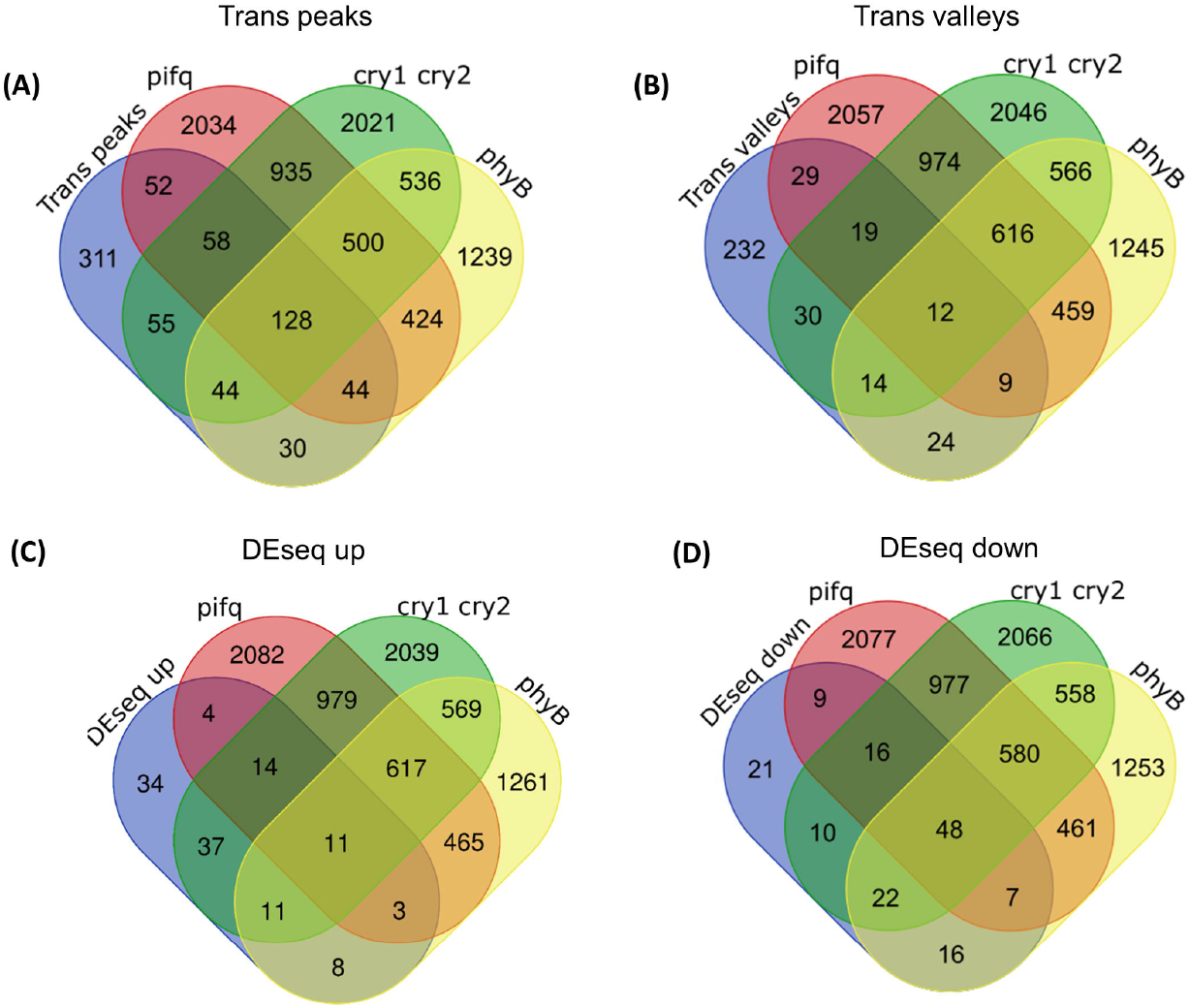
Intersection of commonly regulated genes in shoot light responses for **a** Transcriptome peaks, **b** Transcriptome valleys, **c** DEseq up and **d** DEseq down genes among the five datasets.

### GO term enrichment in transcriptional shoot light responses

The list of significantly enriched GO terms for the genes identified as induced in the Transcriptogram peaks of the three datatsets (GSE5617, GSE29657 and GSE79576) highlights “photosynthesis” followed by “single-organism metabolic process”, “oxidation-reduction process”, “metabolic process”, “cellular metabolic process” along with pigment and cofactor related terms (Fig. 3a), suggesting a strong activation of photosynthesis and photoautotrophy. In addition, significantly overrepresented categories in valleys were related to “drug transmembrane transport”, “peptidyl-proline modification”, “cellular response to auxin stimulus” among others, in agreement with the overall repression of hypocotyl growth triggered by photomorphogenesis in shoots. On the other hand, Diferential Expression (DESeq) upregulated genes were enriched in GO terms “carboxylic acid metabolic process”, “small molecule metabolic process”, “single-organism cellular process”, “single-organism process” and others. The GO terms “single-organism metabolic process”, “oxidation-reduction process”, “cofactor metabolic process”, “small molecule metabolic proces” and “photosynthesis” were the main categories down-regulated in shoot illumination (Fig. 3b). The complete list of GOs terms is listed on Supplementary Table 3. These results suggest that the use of the transcriptogramer seems to have a better coverage related to GO term assessment when comparing different datasets. However, this might be a consequence of the larger number of genes considered.

**Figure 3.**
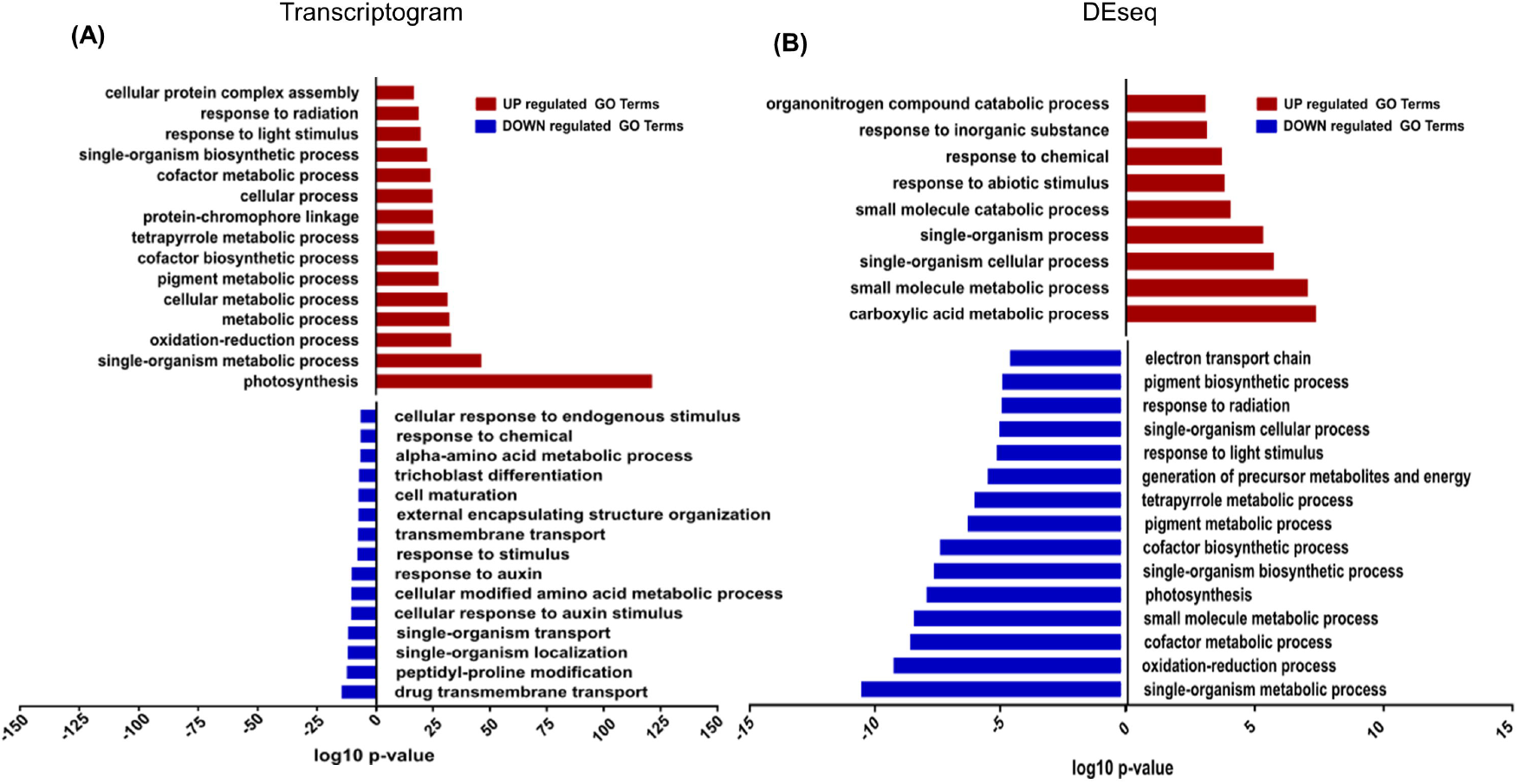
GO term enrichment for light responsive genes from the three datasets (GSE5617, GSE79576 and GSE29657) **a** Functional enrichment analysis of transcriptogram peaks and valleys. The top 15 enriched terms were listed. **b** Functional enrichment analysis of up and down regulated genes identified through DEseq. The top 10 up and 15 downregulated enriched terms were listed. A detailed list of the data is shown in **Supplementary Table 3**.

### Association of light regulated genes with photoreceptor signaling pathways

Light perception by plant photoreceptors CRY1, CRY2, and phytochromes is the main trigger for seedling photomorphogenesis (Podolec and Ulm 2018) which leads to PIF degradation and supression of skotomorphogenesis (Pham et al. 2018). We compared light regulated genes identified by our analysis to differentially expressed genes identified for *pif1pif3pif4pif5* (*pifq*) (Sun et al. 2016), *cry1 cry2* (He et al. 2015) and a contitutively active form of phyB (YHB) (Hu et al. 2009) (Fig. 4). A detailed list of the intersections is presented in Supplementary Table 4. Interestingly, many of the genes identified by DEseq (34 DEseq up and 21 DEseq down) and the majority identified by Transcritogram (311 Peak genes and 232 Valley genes) are not listed as differentially expressed in these light signaling mutant backgrounds neither returned enriched regulators based on cis-regulatory elements by the TF2Network analysis (Kulkarni et al. 2018). However, the majority of the DEseq downregulated (DEseq down) genes (48) were affected in all three mutants (Fig. 4b). The largest single intersection happened with *cry1 cry2*-regulated genes regardless the analysis procedure (DEseq or Transcriptogram), followed by *pifq* and phyB affected genes. The only exception was for DEseq down, where phyB-dependent genes were more represented than *pifq* in the comparison. Although the larger overlap observed with the *cry1 cry2* responsive genes suggests that blue light responses, mediated by cryptochromes, act as an important signal for transcriptional change in early photomorphogenesis, the *pifq-* and phyB-responsive genes belong in a shared signalling pathway where PIF activity is negatively regulated by active phyB (Shin et al. 2009; Leivar et al. 2009). In this context, the majority of differentially expressed genes we identified fall under phyB-PIF regulation.

**Figure 4.**
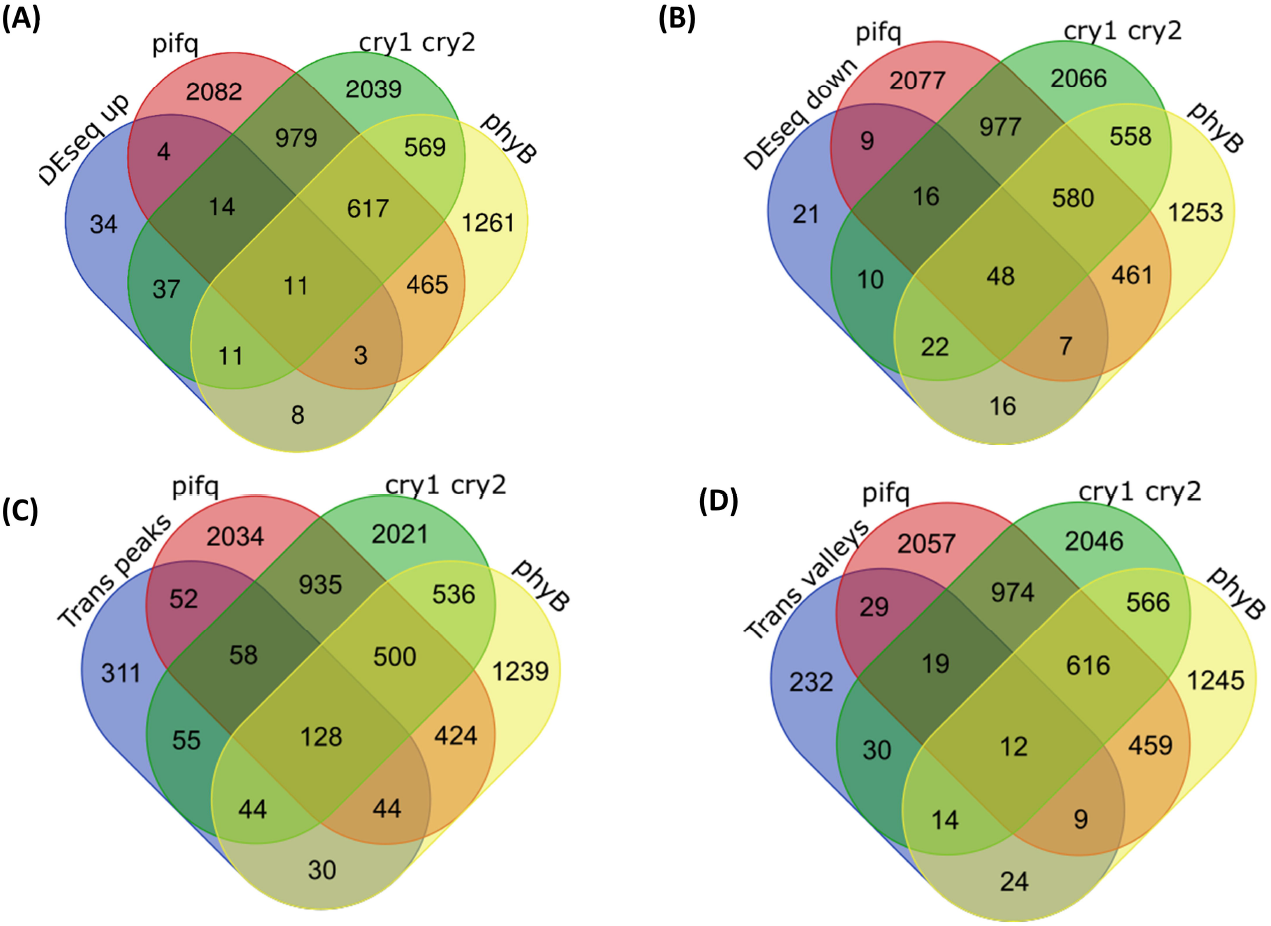
Venn diagram of of light modulated genes from three datasets (GSE5617, GSE79576 and GSE29657) identified by Differential Expression (DESeq) and Transcriptomic Analysis (Trans) overlapping with *pif1pif3pif4pif5* (*pifq*) (Sun et al., 2016), *cry1 cry2* (He et al., 2015) and phyB-YHB (Hu et al., 2009) differentially expressed genes. **a** DEseq up, **b** DEseq down, **c** Trans Peaks, **d** Trans Valleys

The HY5 transcription factor is one of the main promoters of photomorphogenesis; its expression is activated by light and HY5 activates its own expression and many downstram light responsive genes, functioning as a converging hub to many light signals (Podolec and Ulm 2018). So, we also crossed our data with the a refined list of HY5-direct targets (Burko et al. 2020). (Fig. S1, Supplementary Table 5). There was a limited overlap between the genes present in Trans peaks (34), followed by DEseq down (17) and Trans valleys (3). Strikingly, there were no genes in common between the DEseq up and HY5 targets lists (Supplementary Fig. 1a) as one would expect in a list of light-induced genes. This suggests that, the transcriptogram comparison was more robust than DEseq in identifying common light responsive genes and/or, although HY5 is essential to many light responses, it does not seem to act as the main early response transcription factor to the light stimulus, suggesting that other factors might be acting upstream or in paralell to HY5.

### Light regulated gene expression networks overlap with ABA-responsive signaling

During photomorphogenic development, the earliest signaling events downstream of photoreceptor activation do not involve *de novo* protein synthesis. The signaling cascade involves protein stability/activity regulation through protein-protein interactions as well as targeted protein degradation by the 26S proteasome (Hellmann and Estelle 2002). So, transcriptomic analysis alone fails to identify the primary response regulators acting upstream of transcriptional control. In order to identify the putative *cis*-acting regulators of the light responsive genes we identified, we searched their promoter *cis*-elements with the TF2Network (Kulkarni et al. 2018) tool. We selected the top putative regulators based on higher Protein-DNA interaction scores and retreived their known PPI regulatory networks from both DEseq and Transcriptogram gene sets (Fig. 5). Interestingly, the ABA-response bZIP transcription factor ABI5 stood out in both networks (Fig. 5a-c) associated with both up and down regulated genes. Besides, HY5 was present in the DEseq network, reinforcing the possible relationship of HY5 and ABI5 (Bhagat et al. 2021a). ABF1, PIF4, GBF3 and ABF4 were found as major regulators for the DEseq up and Trans peaks and Trans valleys genesets (Fig. 5c). Other putative regulatos identified for the Trans valleys gene set comprise ABF3, GBF2, PIF3, PIF5, CCA1, BES1, AGL9 and AGL20. Putative regulators for the genes exclusively identified for the DEseq up gene set are EIN3, DIV2, LFY and AP3 whereas for the DEseq down only ABI5 showed up as a putative regulator. From this analysis, stands out the abundance of the bZIP-type transcription factors GBFs (G-box binding factor 1-3), the ABFs AREB (ABA Responsive Element Binding Factors 1, 3 and 4) and ABI5 (ABA Insensitive 5) along with bHLH PIFs (PIF3, PIF4 and PIF5). This suggests that modulation of ABA responses mediated by bZIP transcription factors might be a central signaling trigger for photomorphogenesis in *Arabidospsis* shoots.

**Figure 5.**
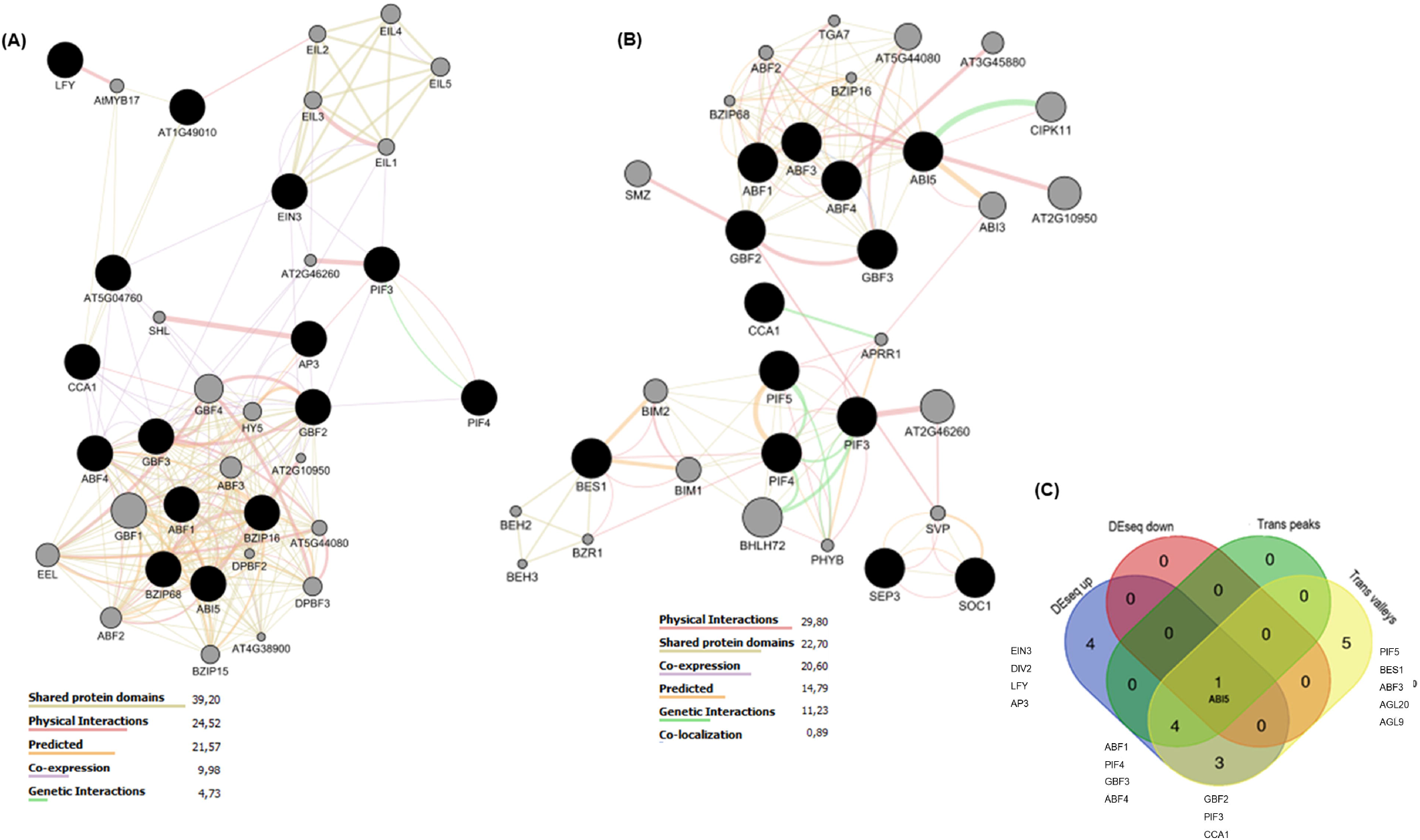
Putative TF networks involved in the regulation of early light responsive gene expression. **a** TF network identified for genes ideintifed by Transcriptogram and **b** Differential Expression (Deseq) analysis. Black circles are transcription factors retrieved from TF2Network and the gray circles represent connections provided by GeneMania. **c** Venn diagram of overlaping putative transcription factors identified for DESeq up and down and Transcriptogram peaks and valleys.

To validate our *in silico* data and to investigate whether the identified transcription factors are involved in light responsiveness, we evaluated the transcriptional response of ABI5 and HY5 genes in some mutants (*abi5-7, abf4, hy5* and *35S:cMYC-ABI5)*. In the dark grown seedlings (0 h), HY5 expression was lower in *abi5-7* and *35S:cMYC-ABI5* when compared to WT (Supplementary table 6). The 1h light-induced HY5 induction observed in the WT, *abi5* and *abf4* was absent ABI5 (*35S:cMYC-ABI5)* overexpression line, similar to the *hy5* mutant (Fig. 6a). Interestingly, the dark (0h) expression in of ABI5 was lower in the *hy5* mutant (Fig. 6b), in contrast with the lower dark expresison of HY5 in *abi5-7* (Fig. 6a). These results indicate that HY5 and ABI5 counteract each other expression in the dark and ABI5 overexpression supresses HY5 induction. It is noticeable the significant decrease of the ABI5 expression in the 35S:cMYC-ABI5 background after transition to light, suggesting that ABI5 RNA might be post-transcriptionally reduced after exposure to light.

**Figure 6.**
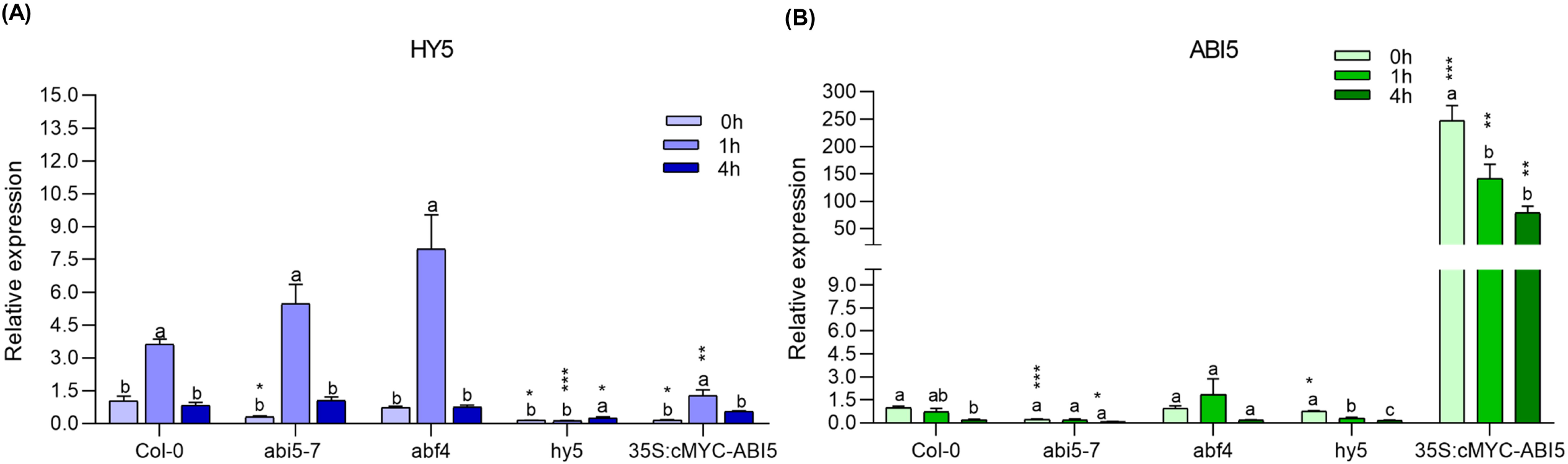
RT-qPCR analysis of gene expression in light treated seedlings. **a** HY5 expression. **b** ABI5 expression. Four-day old dark grown seedlings (0 h) were exposed to white light for 1 (1h) and 4 hours (4h). Statistical significance was determined by ordinary one-way ANOVA with Tukey’s post-test (letters represent the statistical significance between the time within the genotype) and the means were compared by unpaired test t (asterisks denote differentially expressed genes with *p _≤_ 0.05; **p _≤_ 0.01) in the same light condition against the wild-type genotype. Error bars indicate SE.

We evaluated too, the transcriptional response of some putative target genes described as light responsive in seedlings (CHS, CCD4 and NAP). The expression of CHS was more induced after 4 hours of light exposure in the *abi5-7* mutant than in WT (Fig. 7). In the *abf4* mutant, this gene was more expressed in the dark and shows less light inducibility than in WT (Fig.7). For the carotenoid biosynthetic CCD4 gene, *hy5* and *35S:cMYC-ABI5* showed a significant increase after 1 h of light exposure (Fig.7). Interestingly for the NAP gene, all genotypes showed higher expression in the dark when compared to the wild type in this condition. With the exception of *hy5*, the genotypes *abf4, abi5* and *35S:cMyc-ABI*5 showed higher NAP expression than WT in the 4h light treatment.

**Figure 7.**
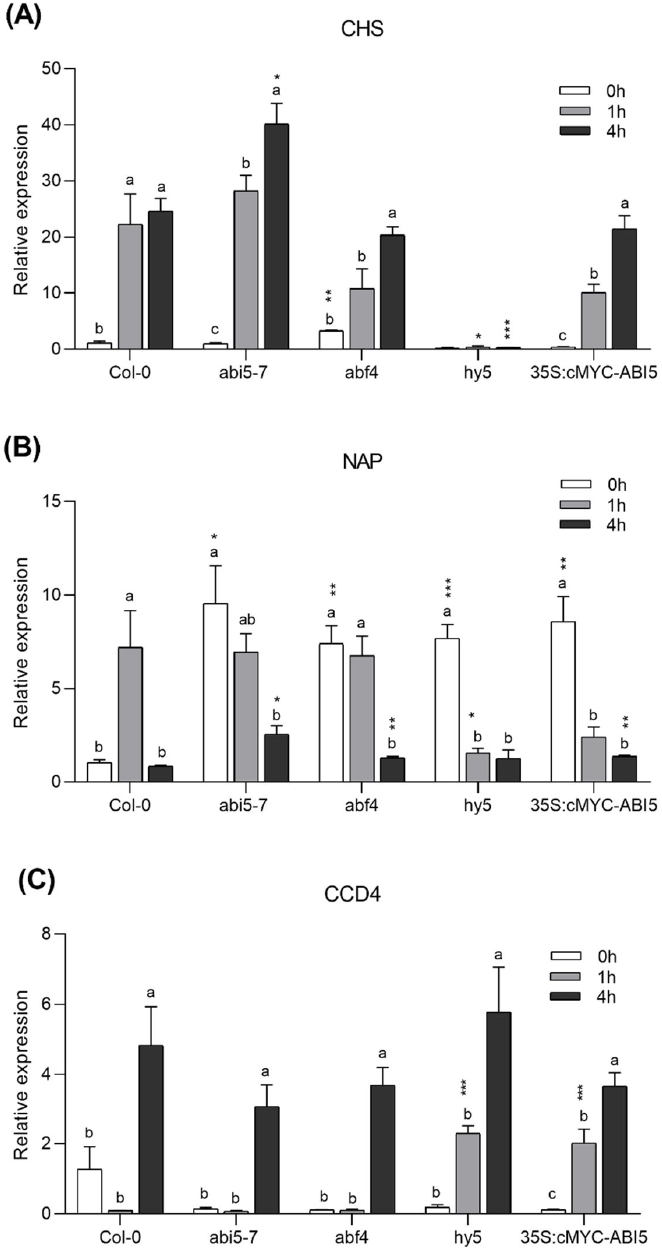
RT-qPCR analysis of gene expression in light treated seedlings. **a** TT4 expression. **b** CCD4 expression **c** NAP expression. Four-day old dark grown seedlings (0 h) were exposed to white light for 1 (1h) and 4 hours (4h). Statistical significance was determined by ordinary one-way ANOVA with Tukey’s post-test (letters represent the statistical significance between the time within the genotype) and the means were compared by unpaired test t (asterisks denote differentially expressed genes with *p _≤_ 0.05; **p _≤_ 0.01) in the same light condition against the wild-type genotype. Error bars indicate SE.

As we observed that the assembly of the photosynmthetic machinery was the top induced process in the trasncriptomic analysis, we decided to evaluate whether the ABA-responsive transcriptional regulators influence this transition. For that purpose, we measured the chlorophyll content during deetiolation in seedlings of WT, *abi5-7, abf3, abf4* and *35S:cMYC-ABI5* exposed to light over 24h (Fig. 8). Total chlorophyll content increased progressively in most genotypes. After 24h of light exposure the *abi5-7, abf3* and *abf4* mutants accumulated almost double the chlorophyll of wild-type suggesting a repressive role over longer light exposures. The *35S:cMYC-ABI5* line had a lower content than the WT after 8-hours of light exposure (Fig. 8a), also suggesting ABI5 has a repressive role in chlorophyll accumulation.

**Figure 8.**
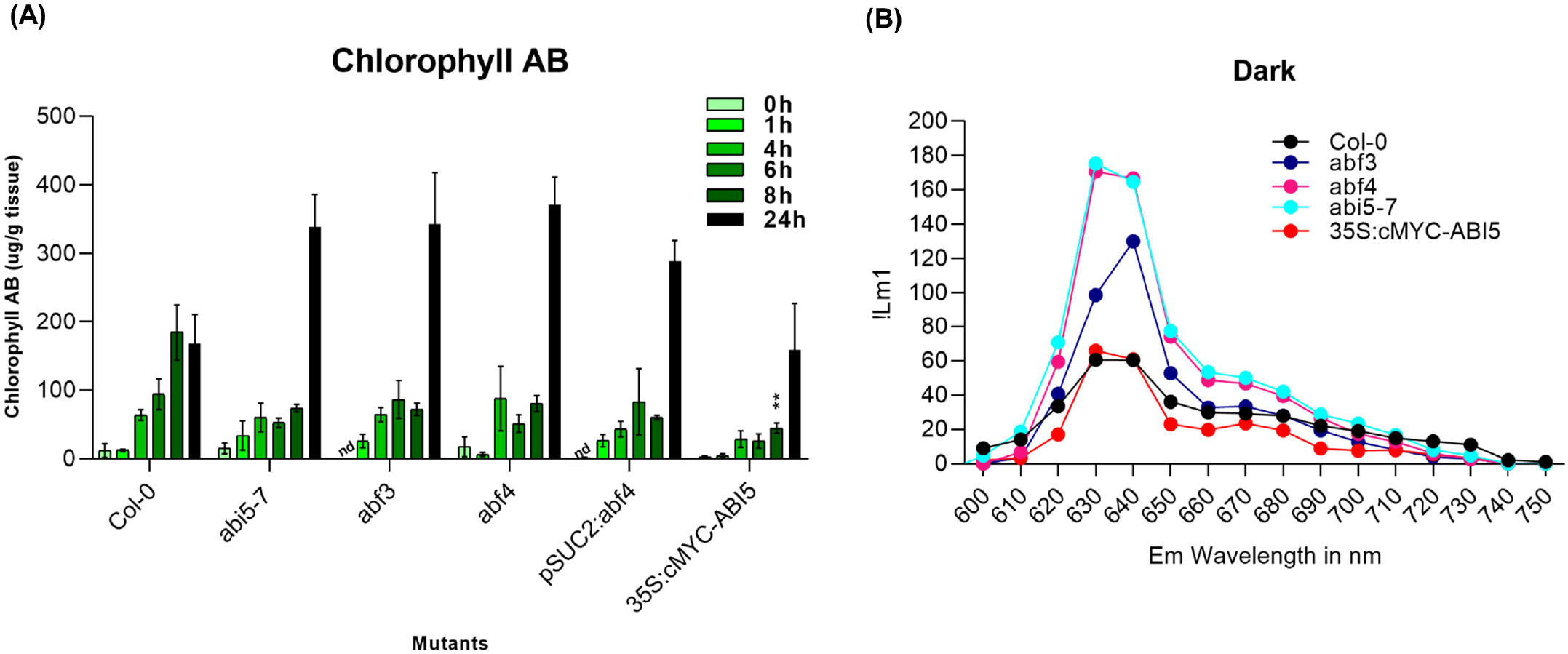
Chlorophyll and protochlorophyllide qauntification in seedlings. **a** Total Chlorophyll (a and b) content in 3-DAG wild-type, *abi5-7, abf3, abf4* and *35S:cMyc-ABI5* seedlings grown in the dark (0h) and after exposure to white light for 1h, 4h, 6h, 8h and 24h.. Error bars indicate SE. n = 3. Asterisks denote significance difference by Kruskal–Wallis test with Dunnett’s post-test (*p _≤_ 0.05; **p _≤_ 0.01) in the same light condition against the wild-type genotype. **b** Fluorescence spectra of acetone extracts indicating protochlorophyllide accumulation of Col-0, *abi5-7, abf3, abf4* and *35S:cMyc-ABI5* seedlings, grown in the dark for 5 d.

The precursor of chlorophyll, protochlorophyllide (Pchlide), is synthesised in the dark and its levels need to be tightly controlled to avoid photobleaching upon illumination. We quantifdied Pchlide levels in 3 DGA dark grown seedlings. Pchlide were higher in *abi5-7* and *abf4*, followed by *abf3* and lower in wild-type and *35S:cMYC-ABI5* (Fig. 8b). These data suggest that these ABA response TFs act in the dark repressing photochlorophyllide production.

### Light controls the spatial expression of ABA response regulators

In order to evaluate the expression pattern of ABA responsive factors during deetiolation, we tested reporter constructs for *pABI5:ABI5-GUS*, pABI5:GUS, pABF4:GUS, 6XABRE_A:GUS. Seedlings were grown for 4 DAG in dark conditions and then exposed to 0, 1, and 4 h light pulses. After 4 day in the dark, *pABI5:ABI5-GUS* expression was detected in the shoot apical meristem (SAM) and cotyledons. After light exposure, the cotyledon expression disappeared whereas the SAM signal remained unaltered showing that ABI5-GUS protein accumulation decreases in cotyledosns. Interestingly, a strikingly different expression pattern was observed for the promoter GUS reporter fusions pABI5:GUS, pABF4:GUS, 6XABRE_A:GUS (Fig. 9). The ABI5 promoter is highly expressed in shoots and roots in the dark, this expression decreases specifically in roots at 1h of light exposure and increasing after 4h again. On the other hand, the pABF4:GUS and the synhtetic ABA response reporter 6xABRE:GUS (Wu 2018) show weak cotyledon and vascular expression and a increasing root expression that is further enhanced by light. These results suggest that the ABI5 protein abundance is repressed by light in the whole seedling during deetiolation except in the SAM. Furthermore, ABF4 and 6xABRE are transcriptionally activated by light in roots possibly to trigger root photomorphogenic development. Therefore, our results suggest that the ABA transcriptional responses are modulated by light, but ABI5 protein accumulation appears not to be quickly repressed by light in the shoot. This suggests that the shoot responses depends of the repreogramming of the ABA response to activate photomorphogenesis.

**Figure 9.**
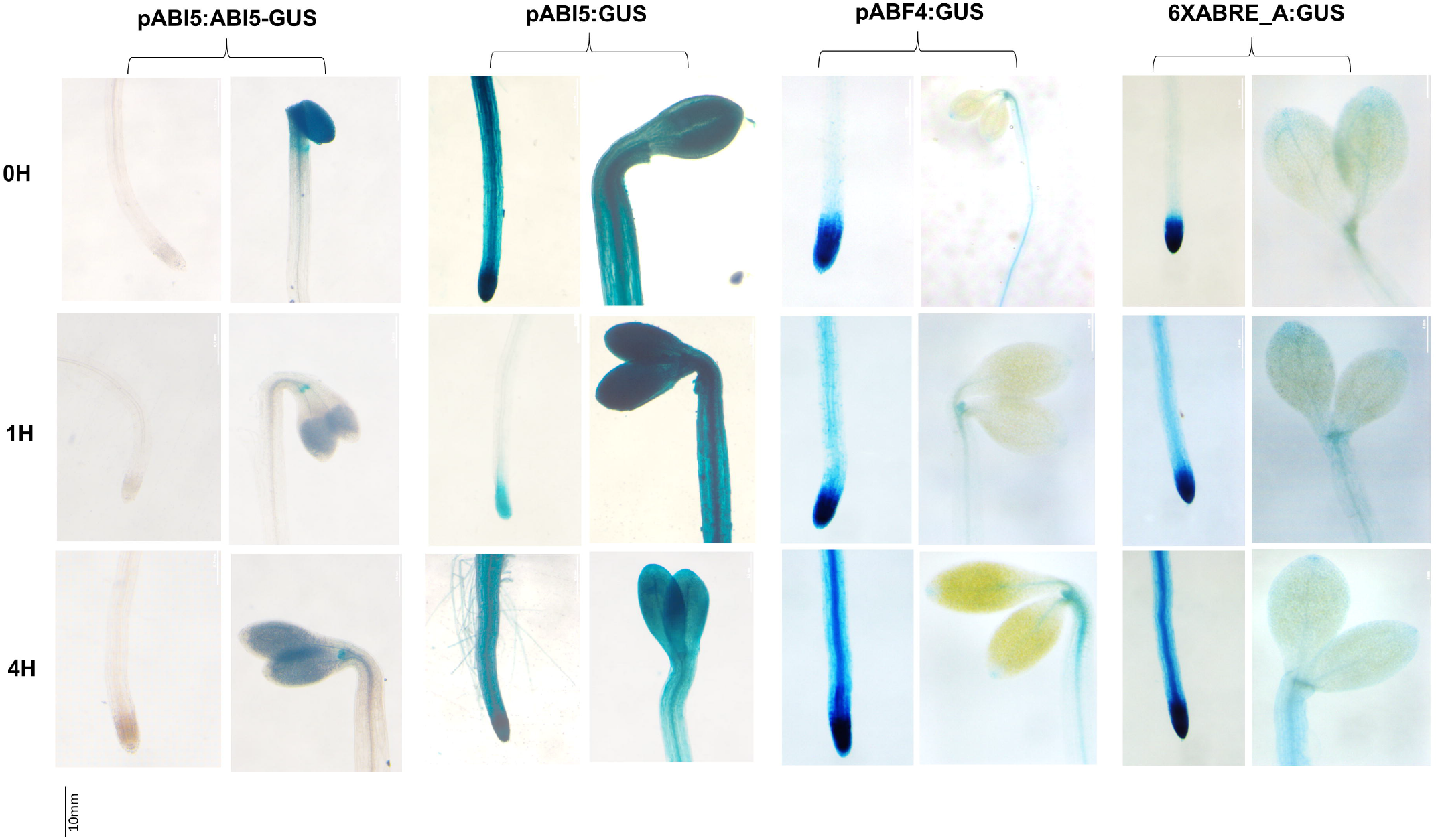
β-glucuronidase assay in *pABI5:ABI5-GUS, pABI5:GUS, pABF4:GU* and *6XABRE_A:GUS* seedlings in darkness (0h), 1h and 4h of white light. Seeds were sterilized and stratified for 3 days at 4°C under dark conditions. Panels represent side by side the primary root and shoots of each timepoint.

### ABA responive regulators are important for seedling photomorphogenesis

In order to evaluate the role of ABA response regulators during seedling photomorphogenic development, we compared the hypocotyl and primary root length of loss-of-function and overexpression genotypes. Both *abi5-7, abf4* and *35S:cMyc-ABI5* showed shorter hypocotyls at 4 DAG similar to the HY5 OE (Fig. 10a), indicating a stronger photomorphogenic response. All tested genotypes had shorter main roots than WT at 10 dag (Fig. 10b). Interestingly, both knockout mutants *abi5* and *hy5* displayed comparable shorter roots (Fig. 10b). These results suggest that ABA responsive factors perform both promoting and repressive effects during seedling photomorphogenesis.

**Figure 10.**
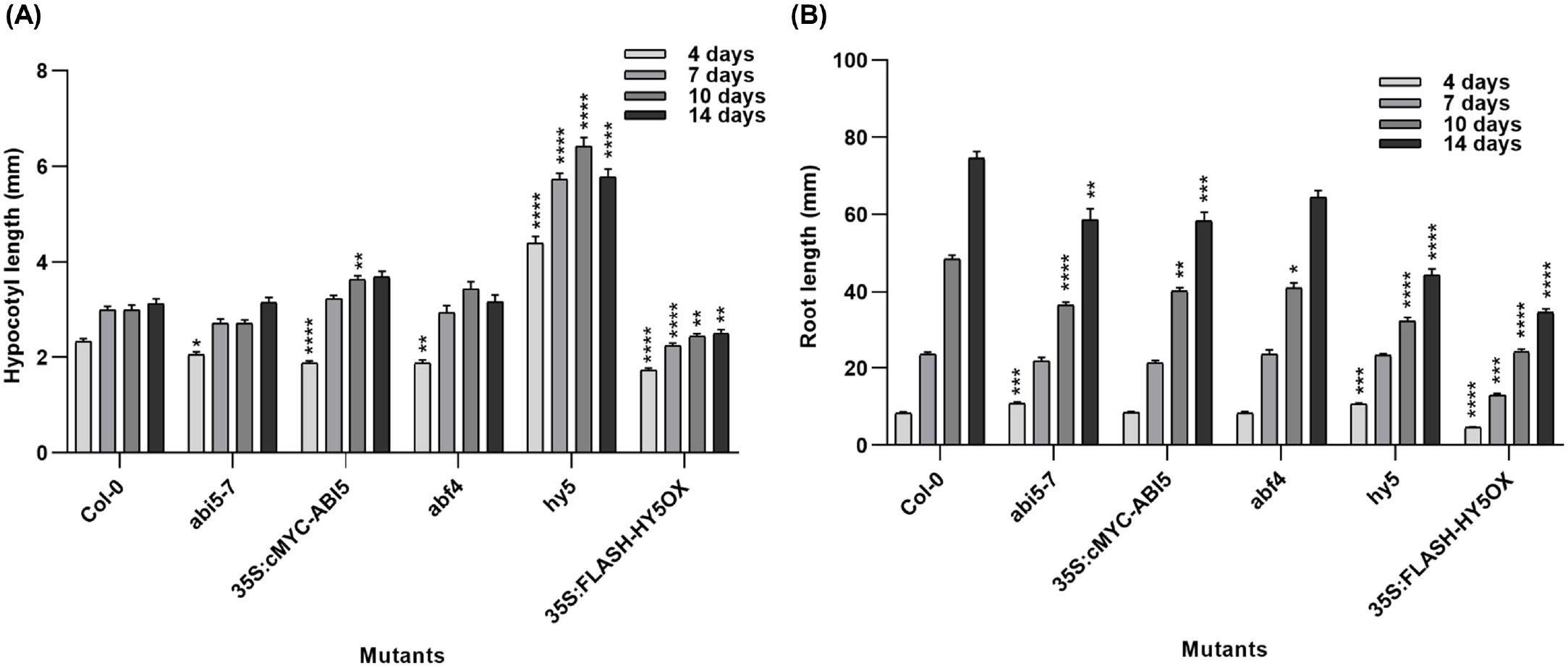
Hypocotyl and root growth lengths of single knockout and overexpression mutants (n ≥ 20) over 14 days of growth. Statistical significance was determined by Kruskal–Wallis test with Dunn’s -test (*p ≤ 0.05; **p < 0.01). Error bars indicate SE.

## Discussion

A large amount of transcriptomic studies have been performed for the model species *Arabidopsis thaliana*. In this work, we integrated data from published transcriptomic databases to identify common regulated genes and process that comprise early shoot light responses. Gene expression profiles from five distinct datasets were examined in order to look for common regulated genes. We report an increase of amplitude of the same transcriptograms peaks and valleys in longer light exposures which suggests that gene expression patterns are sustained qualitatively during photomorphogenesis being quantitatively increased over time (Fig.1b, c). Kilian et al. (2007) (GSE5617) and Miotto et al. (2019) also reported a large number of gene sets in roots affected by shoot□illumination in transcriptogram analysis and a quantitative temporal increase in gene expression patterns in response to light treatment. Hu et al. (2009), comparing the transcriptomes of a constitutive active form of PhyB (YHB) also reported “qualitatively similar to but quantitatively greater” gene expression profiles compared to WT seedlings treated with low red light. These observations suggest that light transcriptional responses are qualitatively specific and quantitatively proportional to the light reatment. Liu et al. 2012 (GSE29657) reported that translational control is central to early photomorphogenesis responses and genes linked to biogenesis of ribosomes and translational machinery are preferentially regulated by translational control, whereas genes related to photosynthetic machinery, chlorophyll and pigments biosytnhesis are more transcriptionally regulated. Nevertheless, once our survey was restricted to transcriptionally regulated processes based on steady state mRNA, our results reinforce his observations. Although we could observe a large amount of similarities from the datasets evaluated, we observed an inversed pattern in the GSE79576 dataset transcript profile related to cotyledon specific samples (light blue line in Figs. 1b, d). One possibility raised by Jiao et al. (2007), is that due to the smaller vacuoles of cotyledons in comparison to hypocotyls, the observed expression patterns might be enriched for the light-induced responses of the apical hook and cotyledons. In the original publication that generated this data, Sun et al., (2016) identified a group of genes (lirSAURs) that were oppositely regulated by light in cotyledons and hypocotyls and these were related to the contrasting light effects in different organs, which may explain this pattern. This observation highlights concerns about evaluting whole seedling expression profiles for cellular processes that might have strong tissue specificities.

Being aware of the restraints imposed by transcritomic analyses, our approach focused on common regulated genes in the different datasets to identify broadly regulated genes for photomorphogenic development. A larger number differentially expressed genes appeared in the Transciptogram analysis compared to Differential Expression. This is due to the fact that the Transciptogram method genes due to the “neighborhood” between them. The transcriptogram is a method that encompasses the entire genome to analyze genetic expression data, it calculates the average expression of a set of genes with similar functions, allowing a global view of the metabolic pathways which are coordinately regulated. It consists of an one-dimensional rearrangement of the genome based on protein-protein associations (Franceschini et al. 2012). This reordering clusters genes associated with the same pathways, allowing to globally compare if a pathway is repressed or induced (Rybarczyk-Filho et al. 2011; da Silva et al. 2014). Functional enrichment demonstrated that the categories of GOs terms identified for the DEseq gene set were not very informative (Fig. 3b) when compared to those obtained for the Transcriotogram (Fig. 3a) where terms “Photosynhtesis”, “Tetrapyrrole metabolic process”, “Protein chromophore linkage” and “Pigment metabolic processes” were over-represented in the induced peaks. The GO terms of the Trans valleys indicate the repression of “Peptidyl-proline modification” which is an essential modification for the expansion of the cell wall (Zdanio et al. 2020) and “response to auxin” which is essential for hypocotyl elongation (Du et al. 2022) (Fig. 3b), are more logical than the unespected list of repressed processes we identified by DEseq terms which find “Photosynthesis” and “Pigment metabolic processes”, “Response to light stimulus” as putatively down regulated by light. Our results with the Transcriptogram tool showed enrichment terms not identified by DEseq, reinforcing the idea that different approaches to data analysis can lead to different sets of genes and processes. From our point of view, for comparison of different datasets, the Transcriptogram appears to select for more robust expression patterns, allowing better identification of core commonly regulated processes than DEseq.

Photomorphogenic responses are dependent of photoreceptor activation, mostly through the action phytochromes and cryptochromes repressing COP1 and PIFs activities. Previous works have identified transcriptomic responses that are altered in *pifq, cry1cry2* and *phyb(YHB)* mutants, allowing us to evaluate which signaling components are required for the light signaling that regulated these datasets. We found that many light-regulated genes belong in the *cry1cry2*-dependent dataset, suggesting that blue light, acting through cryptochromes plays a central role as a trigger for early photomorphogenic transcriptional control. Although cryptochrome activation represses hypocotyl growth, the most represented GO categories in CRY1-regulated genes (He et al. 2015) are “Photosynthesis” and “Chloroplast organization” followed by growth related terms such as “Cell wal organization” and “Hormonal responses”, reinforcing that the establishment of the photosynthetic machinery is the main transcriptionally activated process in early photomorphogenesis. Cryptochromes also mediate blue-light dependent cell expansion in cotyledons and chloroplast development (Yu et al. 2010). This allows identification of the hypocotyl and cotyledon blue light responses which were identified in our datasets. CRY1, along with phyB, acts on repression of auxin signaling through light-induced interation with AuxIAA proteins to inhibit hypocotyl elongation (Xu et al. 2017). We found GO terms “Cellular response to auxin stimulus” and “Response to auxin” enriched in the Transcriptogram valleys of light repressed genes, which associated the supression of auxin response downstream of photoreceptors as a central photomorphogenic response.

Phytochrome activation is also central to photomorphogenic transcriptional responses to red/far-red light (Tepperman et al. 2001; Tepperman et al. 2004; Quail 2007). Activated phytochromes modulate gene expression through direct interaction with nuclear localized PIFs leading to their proteasomal destruction and supression of dark-induced genes (Al-Sady et al. 2006; Leivar et al. 2009; Pham 2018) as well as supression of COP1-SPA complexes by protein-protein interactions (Sheerin et al. 2015; Zhu et al. 2015). We found the largest overlap of light modulated genes in common with *pifq* (Sun et al., 2016), and *phyB* (Hu et al. 2009) affected genes. This observation is consistent with the major role of the PhyB-PIF signaling pathway in controlling early seedling light transcriptional responses (Shin et al. 2009; Leivar et al. 2009; Zhang et al. 2013). PIFs are important repressors of light responses (Shin et al. 2009) and their light-dependent degradation is essential for the initial steps of photomorphogenesis. The *pifq* mutant displays a constitutive photomorphogenic phenotype in the dark (Leivar et al. 2008), with a high expression of chlorophyll biosynthetic genes and photosynthetic apparatus proteins, similar to plants overexpressing a constitutively active form of phyB (YHB) (Hu et al. 2009).

By analyzing the transcription factors binding sites overrepresented in the promoters of the light responsive genes identified in our analysis, we were surprised not to find classical photomorphogenic activators such as HY5 or HYH (Oyama et al. 1997; Holm et al. 2002) but many to be involved with Abscisic Acid (ABA) response. The ABA-responsive A-type bZIPs (ABFs, ABI5) and G-type bZIPs (GBF2 and GBF3) were all centrally located in the PPI networks of putative TFs controlling dark-to-light transcriptional changes. Plant bZIP proteins preferentially bind to the *cis*-elements with a ACGT core sequence, such as G-box (CACGTG), C-box (GACGTC), Z-box (ATACGTGT), and A-box (TACGTA). The Transcriptogram network (Fig. 5b, c) also linked members of the PIF family (PIF3, PIF4 and PIF5) in a cooperative network. PIFs are activators of the ABA response, which is usually induced in the dark in cotyledons, and are degraded in light. It has been reported that much of the photomorphogenic response involves suppressing the ABA response via PIFs and phyB (Ha et al. 2018). PIF1 interacts physically with ABI5 and other A-type bZIP transcription factors to enhance PIF1 binding to a set of target sites in vivo (Kim et al. 2016). PIF4 activates ABI5 expression in the dark (Qi et al. 2020). Part of the PIFs control of seedling growth, involves ABA signaling (Qi et al. 2020; Liang et al. 2020). The G-box binding factors (GBF1-3), also appeared largely connected to the PPI network (Fig. 5a, b, c). GBFs are ABA-responsive genes regulated by PIFs (Oh et al. 2009b). GBF1 is a blue-light responsive TF acting downstream of the cryptochromes controlling blue-light specific responses. GBF1 overexpression reduces blue-light repression on hypocotyl elongation (Mallappa et al. 2006). GBF1 is degraded in the dark by the proteasome in a COP1-independent manner, and GBF1-COP1 interaction stabilizes GBF1 in the light (Mallappa et al. 2008b). GBF1 interatcs with both HY5 and HY5 balancing their regulatory effects (Singh et al. 2012). On the other hand, GBF3 is more abundantly expresed in darkness and roots, and it can heterodimerize with other GBFs (Schindler et al. 1992). GBFs share many co-regulated genes with HY5 and HYH and change their interaction partners depending on the tissue and light. Although some are constitutively expressed (GBF1 and GBF2), their partners can change reprogramming the developmental output (Schindler et al. 1992; Kurihara et al. 2020). We suggest a scenario were, in the dark and absence of HY5, GBFs control a set of dark-response genes and, upon illumination, GBFs can physically interact with HY5 which accumulates and displaces GBFs to different promoters activating photomorphogenesis (Singh et al. 2012).

ABA INSENSITIVE 5 (ABI5) was the only common transcription factor putatively regulating the four sets of genes (Fig. 5a, b, c) overlapping with induced and represses gene sets. ABI5 is a positive regulator of ABA signaling, expressed in cotyledons, hypocotyls, and roots of young seedlings. ABI5 is involved in postgermination developmental arrest, repression of germination and early seedling development. ABI5 regulates genes both during seed development, germination and post germination growth (Skubacz et al. 2016). ABI5 enhances the binding of PIF1 to target promoters to inhibit seed germination (Cheng et al. 2014; Kim et al. 2016). Several previous studies have identified factors that promote photochlorophilide accumulation in the dark as *pif1* and *pif3* mutants have higher levels of photochlorophilide (Job and Datta 2021). Our studies show that ABI5, ABF3 and ABF4 can act as potential repressors, since their mutants showed a higher levels of photochlorophilide in the dark compared to wild type (Fig. 8b).

ABI5 is positively regulated by HY5 (Chen et al. 2008) and also promotes its own expression which represents an integration light and ABA signaling process (Xu et al. 2014; Jing and Lin 2020). It has been identified that HY5 and ABI5 interact and integrate light and ABA responses, indicating that ABI5 can act as a negative regulator in photomorphogenesis and HY5 acts as a positive regulator of ABA signaling (Bhagat et al. 2021). The overexpression of ABI5 inhibits hypocotyl elongation under blue, red, or far-red light in Col-0 (Chen et al. 2008; Xu et al. 2014), which places ABI5 as a component of photomorphogenic responses. Based on our observations that the majority of early light transcriptional responses might depend on ABA response factors which are already present in the dark grown seedling, we propose a model where light signaling repurposes part of this available transcriptional machinery to activate the early responsive genes upon PIF degradation and later on, the stabilization of positive regulators of light responses (e.g. HY5, HYH, LAF1, GBFs, among others) takes over to sustain photomorphogenic growth (Fig.11). This effect might occur by direct interaction with ABA-responsive bZIPs, as HY5 and ABI5 have recently been found to physically interact (Bhagat et al. 2021). Other recently study demostrated that, COP1 modulates ABA signaling during seedling growth in dark condictions by regulating ABA-induced ABI5 accumulation, suggesting that plants adjust their signaling machinery to the ABA according to luminosity in which they find themselves (Du et al. 2022). In summary, through a survey of transcriptomic datasets, the transcriptogram method allowed us to identify common light regulated processes central to early seedling photomorphogenesis and propose a novel transitory transcription cascade for deetiolation dependent on ABA-responsive bZIP trancription factors guiding photomorphogenic development.

**Figure 11.**
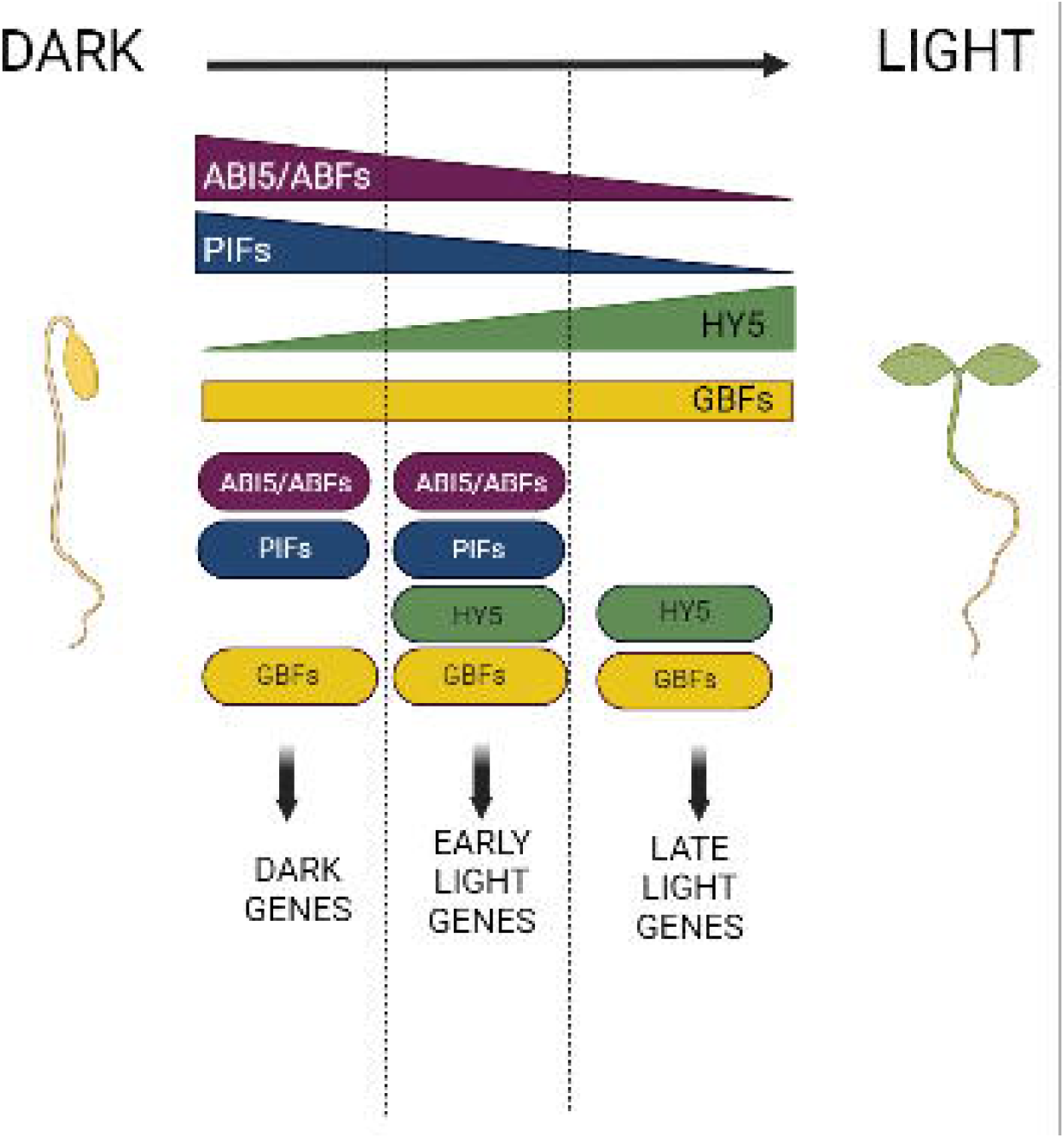
Proposed working model for the activation of early photomorphogenesis. PIFs, ABI5/ABFs and GBFs regulate the expression of skotomorphogenic genes the dark (Dark Genes) where PIFs and ABI5/ABFs act cooperatively to regulate ABA responses. ABI5 and PIFs protein abundance is progressively reduced after exposure to light whereas HY5 abundance increases. During the induction of early light responsive genes, the decaying transcription factors ABI5/ABFs and PIFs interact and cooperate with light responsive regulators GBFs and HY5 to initiate photopmorphogenesis (Early Light Genes). GBFs are continuosly expressed. After prolonged light exposure ABI5/ABFs and PIFs are degraded and positive photomorphogenic regulatos overtake transcriptional control (Late Light Genes). Figure created using Biorender.

## Acknowledgements

We thank Dr. Fernanda Lazzarotto for the critical reviewing of the manuscript. This study was financed in part by Conselho Nacional de Desenvolvimento Cientifico e Tecnológico (CNPq) and Coordenação de Aperfeiçoamento de Pessoal de Nível Superior – Brasil (CAPES) Finance Code 88881.068110/2014-01.

## Author contributions

CFS— analyzed the data, wrote the manuscript. RJSD and IS—processed the sequencing raw data. BHO and LA—helped with the transcriptogram analysis. FSM—conceived, supervised the study and wrote and edited the manuscript. All the contributors read and approved the final version.

## Figure captions

**Supplementary Figure 1** Venn diagram of genes from Differential Expression (DESeq) **a** up, **b** down and Transcriptomic Analysis (Trans) **c** peaks, **d** valleys, from three datasets (GSE5617, GSE79576 and GSE29657) of light modulated genes and HY5 target genes (Burko et al., 2020)

**Supplementary Figure 2** *Arabidopsis thaliana* transcriptogram of dataset GSE79576. The x-axis relative to gene position have been divided by the total number of proteins retrieved from STRING. Average transcriptograms for r=90, signifcant peaks (up-regulation) or valleys (down-regulation) are colored in blue. Grey lines show non-signifcant regions.

**Supplementary Figure 3** *Arabidopsis thaliana* transcriptogram of dataset GSE29657 and GSE5917. The x-axis relative to gene position have been divided by the total number of proteins retrieved from STRING. Average transcriptograms for r=90, signifcant peaks (up-regulation) or valleys (down-regulation) are colored in blue. Grey lines show non-signifcant regions.

**Supplementary Figure 4** RT-qPCR analysis of gene expression of chlorophyll related genes in seedlings. Gene expression was normalized using an internal control (; Miotto et al., 2019) for each reaction. Statistical significance was determined by ordinary one-way ANOVA with Tukey’s post-test (letters represent the statistical significance between the time within the genotype) and the means were compared by unpaired test t (asterisks denote differentially expressed genes with *p _≤_ 0.05; **p _≤_ 0.01) in the same light condition against the wild-type genotype. Seedlings were grown under 94 _μ_mol m _− 2_ s _– 1_. Error bars indicate SE.

**Supplementary Table 1**. Datasets used in the meta-analysis

**Supplementary Table 2** Differentially expressed genes and genes present in peaks or valleys of transcriptogram, in common between the three datasets (GSE79576, GSE28657 and GSE5617) in their conditions.

**Supplementary Table 3** Functional enrichment analysis of list of genes present in DEseq up or down and peaks or valleys from three datasets select (GSE5617, GSE79576 and GSE29657). Bold terms were those plotted Fig. 3.

**Supplementary Table 4** Differentially expressed genes and genes present in peaks or valleys of transcriptogram, in common between the three datasets (GSE79576, GSE28657 and GSE5617), with expression value in papers associated the *pifq* (Sun et al., 2016), *cry1 cry2* (He et al., 2015) and phyB-YHB (Hu et al. 2009).

**Supplementary Table 5** Differentially expressed genes in common between the three datasets (GSE79576, GSE28657 and GSE5617) compared to HY5 targets (Burko *et al*., 2020).

**Supplementary Table 6** RT-qPCR primers. List of RT-qPCR primers used for gene expression analysis of Figure 7.

