## Supplementary figures and images for "A survey of transcriptomic datasets identifies ABA-responsive factors as regulators of photomorphogenesis in *Arabidopsis*"

### Suplementary Figure 1

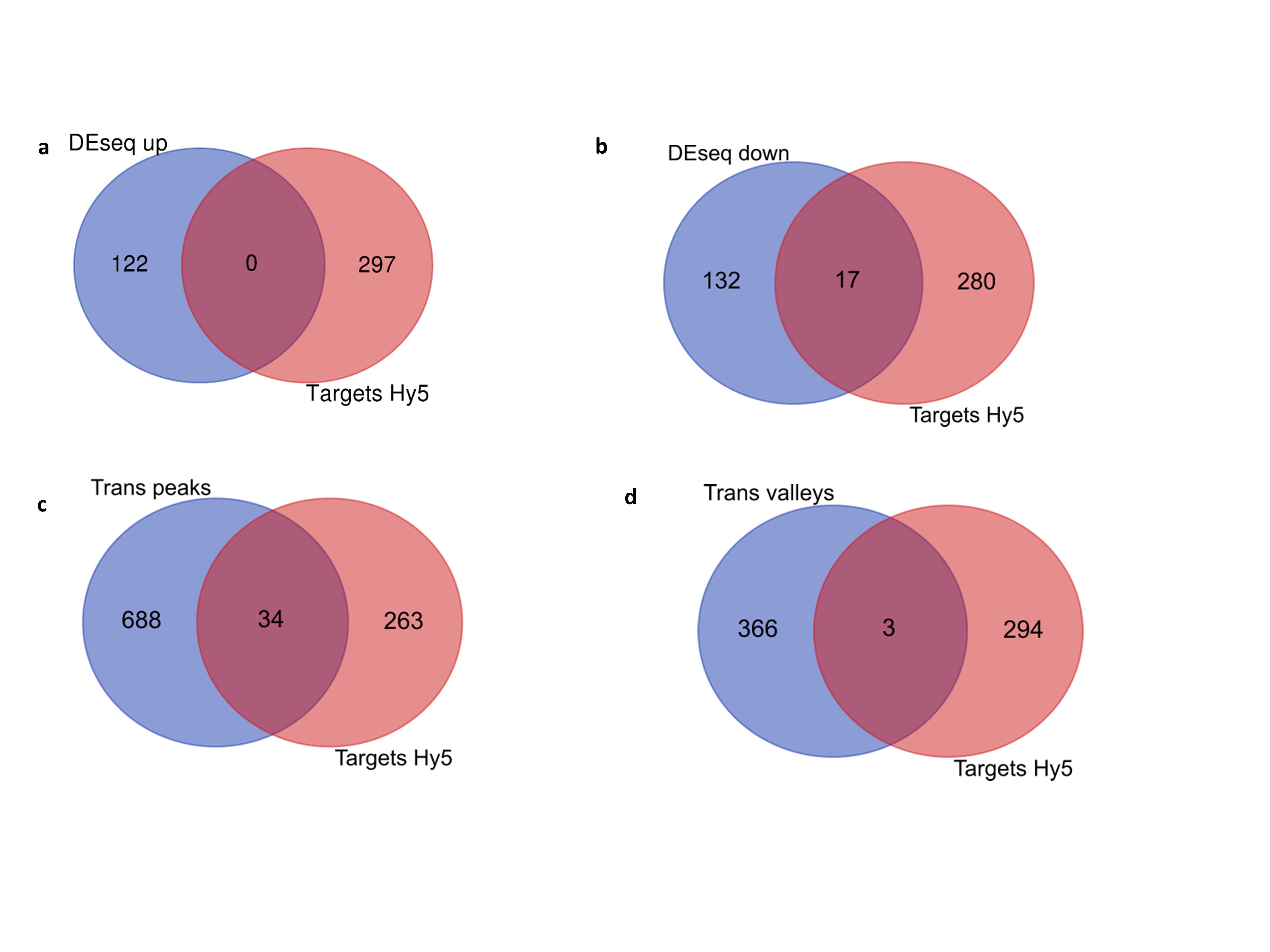

### Suplementary Figure 2

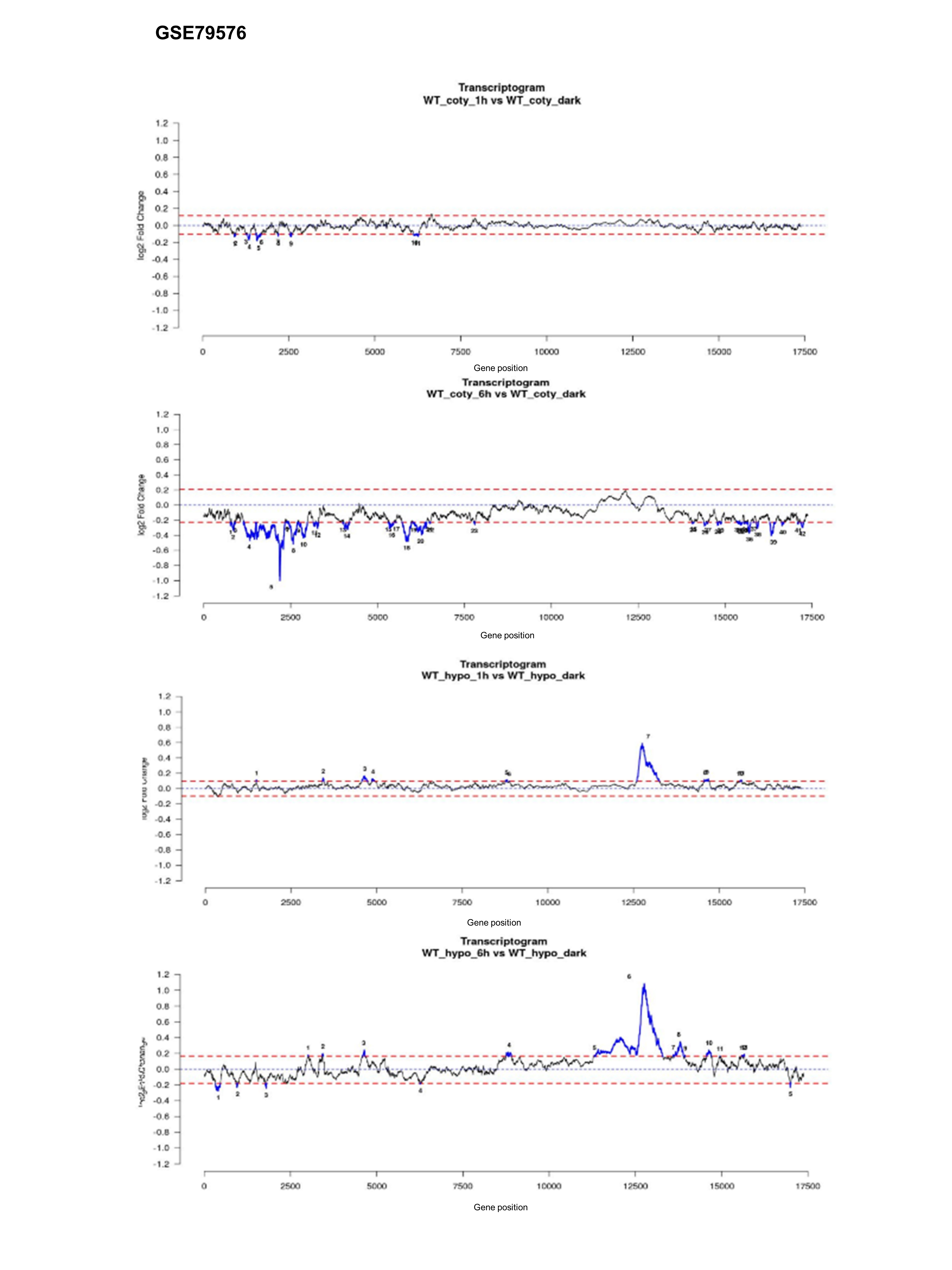

### Suplementary Figure 3

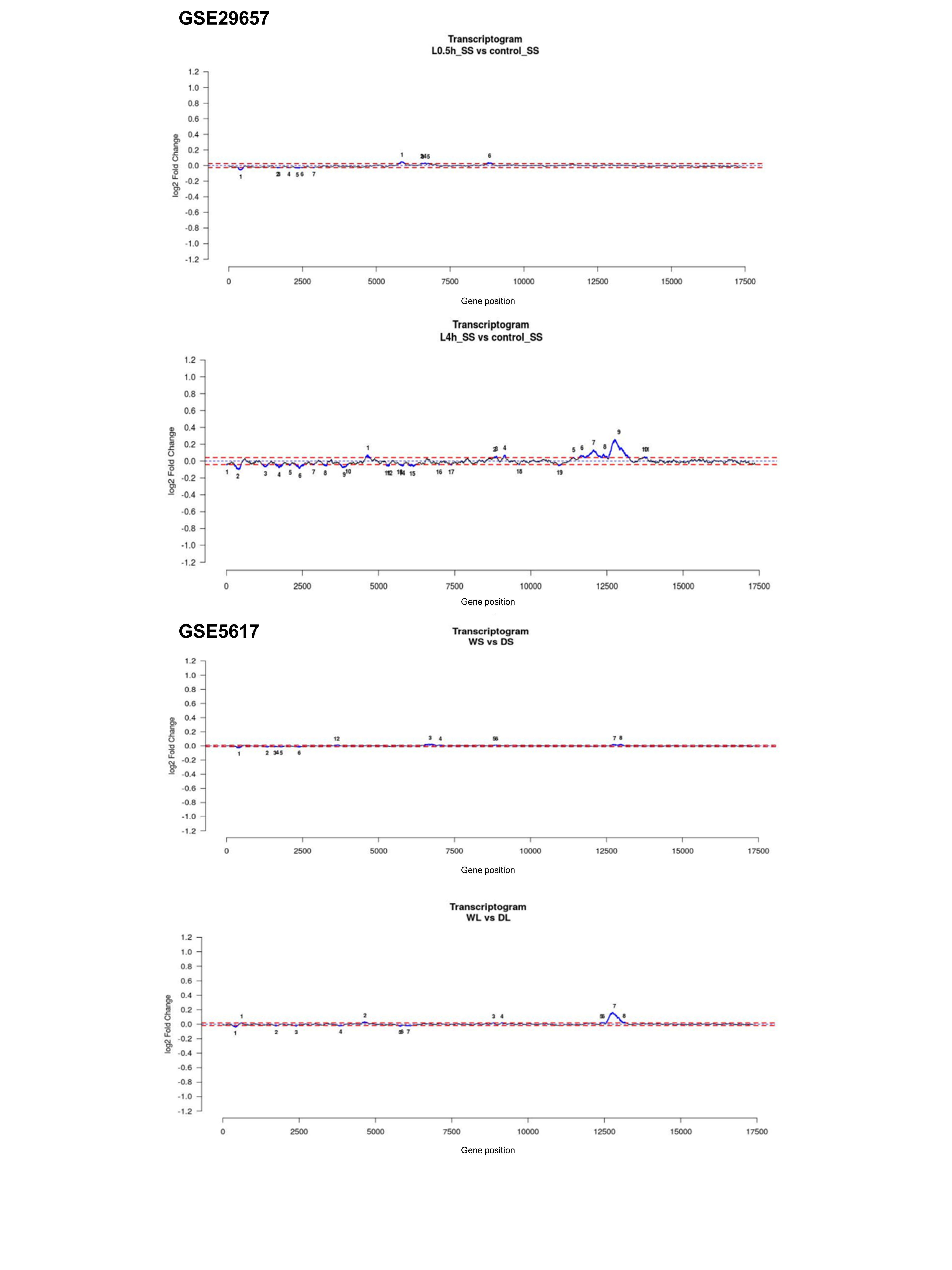

### Suplementary Figure 14

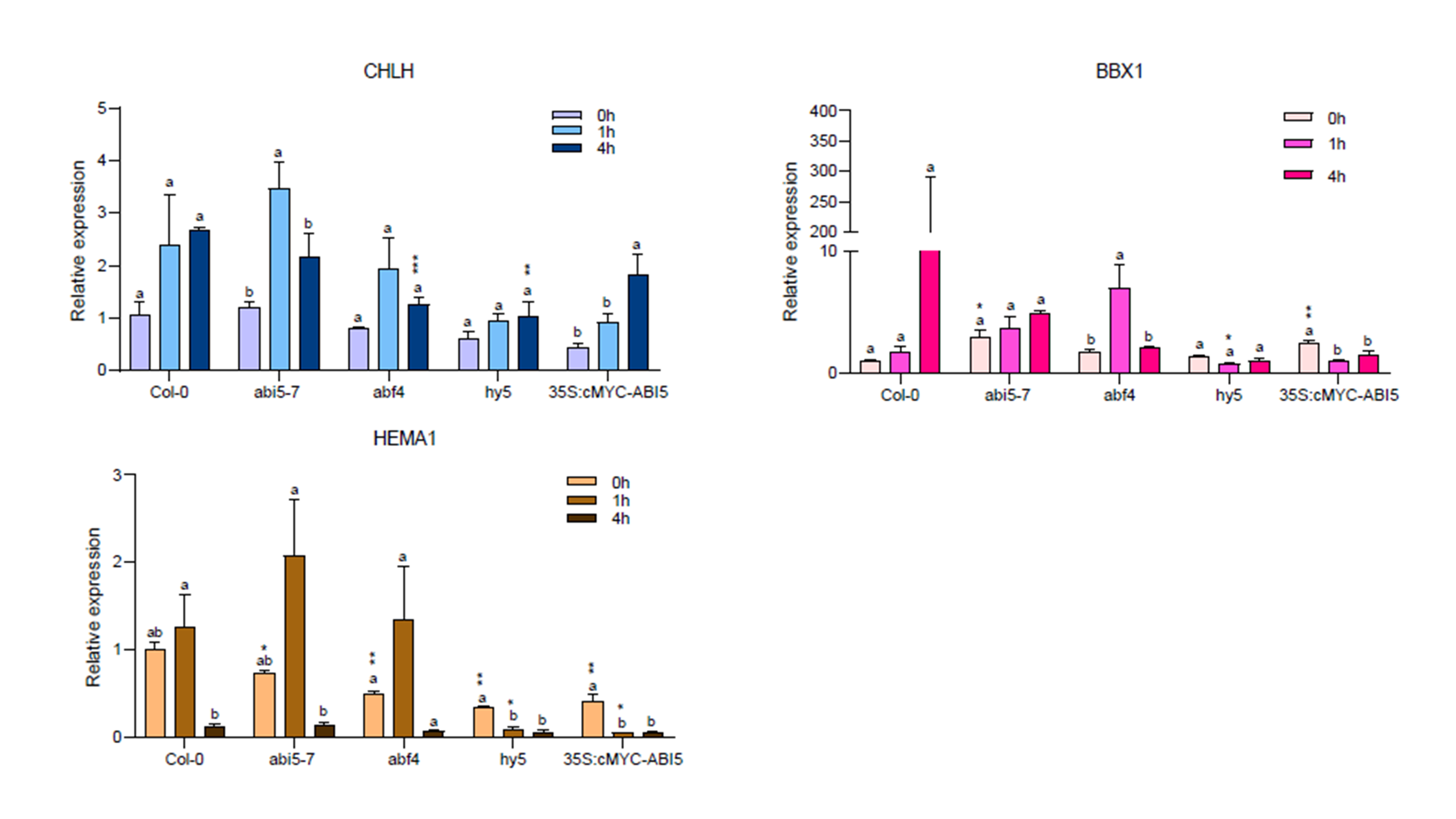
